# Experience shapes initial exploration for non-generalizable spatial learning

**DOI:** 10.1101/2023.12.26.573368

**Authors:** Michelle P. Awh, Kenneth W. Latimer, Nan Zhou, Zachary M. Leveroni, Zoe M. Stephens, Jai Y. Yu

## Abstract

Experience can change how individuals learn. Learning to solve a new problem can be accelerated by generalizing known rules in the new context, but the impact of experience on solving problems where generalization cannot be applied remains unclear. To study the impact of experience on solving new problems that are distinct from previously learned tasks, we examined how rats learned a new spatial navigation task after having previously learned different sets of spatial navigation tasks. The new task differed from the previous tasks in spatial layout and navigation rule, and could not be solved by applying previously learned rules. We found that different experience histories did not impact task performance in the new task. However, by examining navigation choices made by rats, we found exploration patterns during the early stage of learning in the new task was dependent on experience history. We identified these behavioral differences by analyzing each rat’s navigation choices and by modeling their choice sequences with a modified distance dependent Chinese restaurant process. We further pinpointed the behavioral difference to sequential turn/no turn decisions made at choice points. Our results indicate that experience can influence problem-solving strategies when learning to solve new problems. Individuals with distinct experience histories can approach new problems from different starting points but converge on the same solution.

## Introduction

Individuals learn from distinct and diverse experiences to build general knowledge, which can be applied to new problems (Alonso et al., 2020). New problems can be solved faster when aspects of previous experience, such as learned rules, can be directly applied (Thorndike and Woodworth, 1901a; Harlow, 1949). When the new problem is distinct from previous experience, the effect of past experience becomes challenging to understand since improvements in performance on these problems are less consistent (Thorndike and Woodworth, 1901b; Wiltbank, 1919).

Two lines of research provide different perspectives. At the cognitive level, diverse experiences are thought to improve general problem-solving ability. Experiential diversity in the form of environmental enrichment has revealed neuroanatomical (Bennett et al., 1964; Diamond et al., 1964; Heller et al., 2020; Urban-Wojcik et al., 2021; Bogado Lopes et al., 2023) and behavioral changes (Leggio et al., 2005; Nithianantharajah and Hannan, 2006; Petrosini et al., 2009; Freund et al., 2013; Gelfo, 2019) in humans and animals. These findings support the hypothesis that having diverse interactions with the environment leads to brain-wide changes that are not linked to specific experiences. Alternatively, studies in the early 20^th^ century found rats, non-human primates, and humans could solve new problems faster after previously encountering related but not identical problems (Thorndike and Woodworth, 1901a; Thorndike and Woodworth, 1901c; Thorndike and Woodworth, 1901b; Wiltbank, 1919; Ho, 1928; Thorndike, 1935; Harlow, 1949).

A proposed explanation for this effect is “transfer” of learning, which is generally quantified by changes in performance metrics, such as the number of attempts to reach criterion, the number of errors made, or the time needed to solve a task. A range of transfer outcomes have been observed, ranging from positive transfer (improvement in performance) to neutral effect (no change) to negative transfer (worsening) (Webb, 1917; Wiltbank, 1919; Dennis et al., 1932). Further, transfer was reported for different degrees of experience: rats performed fewer errors when learning a new maze even with partial training of a different maze (Ho, 1928; Bunch and Lang, 1939) or after training in multiple mazes (Dashiell, 1920; Rashid et al., 2017). Further, solving new problems can also improve when rats had dissimilar past experience, such as prior operant learning improving new spatial learning (Adams, 2003) or positive transfer between mazes with different rules (Gallup and Diamond, 1960). Thus, previous experience has complex effects on future problem-solving. Although these experiments identified differences in performance in new tasks depending on the experience of the animals, it is unclear how aspects of learning behavior were affected. Specifically, does the absence of transfer, as defined by a lack of performance changes, indicate previous experience had no effect on future learning?

Performance metrics across time, such as speed or accuracy, are used in these studies to determine the impact of past experience on solving new problems. Reductions in the numbers of errors or in the time or trials taken to solve the task were taken as evidence of the transfer of previous knowledge to the solving of novel tasks. In experiments involving learning multiple tasks over time, improvements in performance metrics across consecutive learning sessions were not always observed (Wiltbank, 1919; Dennis et al., 1932). On one hand, this could indicate a lack of consistent transfer or generalization between some tasks. Alternatively, the metrics used to quantify performance may have failed to capture aspects of behavior that did differ. This is especially relevant to exploration in structured environments since humans and non-human animals show intricate exploratory patterns (Uster et al., 1976; Alonso et al., 2021; Rosenberg et al., 2021; Brunec et al., 2023). Thus, examining detailed aspects of behavioral choices in addition to traditional performance metrics can provide important insight on how learning occurs.

Here, we investigate how past experience affects the way animals learn a new task that could not be solved by directly applying previously learned rules. We compare spatial learning in a novel task between groups of rats that previously learned either one or two spatial tasks that differed in topology and rule. We did not find experience-dependent differences in performance based on reward rate over time. However, we reasoned that differences could lie in strategic choices during exploration that may not be reflected in gross performance measures. To search for additional experience-dependent influences on learning, we quantified patterns of spatial decisions during exploration. Surprisingly, we found experience-dependent differences in these spatial exploration patterns only during initial learning. These differences disappeared after rats discovered the rule and their behavior converged in the new spatial task. Our results show experience alters the starting state of animals, affecting how they discover rules for unfamiliar problems, and this can occur without affecting learning performance.

## Results

To understand how learning a novel task depends on previous experience, we designed a two-phase spatial learning experiment. The first phase of the experiment is the differential experience phase, where rats were assigned to “diverse” or “uniform” experience conditions (Fig. 1A). In the diverse experience groups (n=9), rats were trained on two mazes each day, the H maze (H) or the double T maze (2T), counterbalanced for session order (n=5 and n=4). During training, the rat could explore all arms of the mazes, but the task rule required rats to visit two specified arm ends in alternation to receive reward. Uniform group rats (n=10) were trained on only the H or 2T maze, twice per day (Fig. 1A) (n=5 for each maze). To create distinct spatial navigation experiences that took similar physical effort, the H and 2T mazes differed in geometry but shared the same topology. The mazes each featured two intersections, and the rewarded trajectories had the same path lengths. After training, all animals achieved similar levels of performance in their Phase 1 tasks (Fig. 1C). The second phase of the experiment is the common experience phase, where all animals were given the same new spatial task: a Plus maze with an alternation rule (Fig. 1B) for two sessions per day over 5 days. The new task differed from the H and 2T tasks in topology, geometry, and reward rule, and thus could not be solved by applying spatial navigation rules from phase 1.

**Figure 1.**
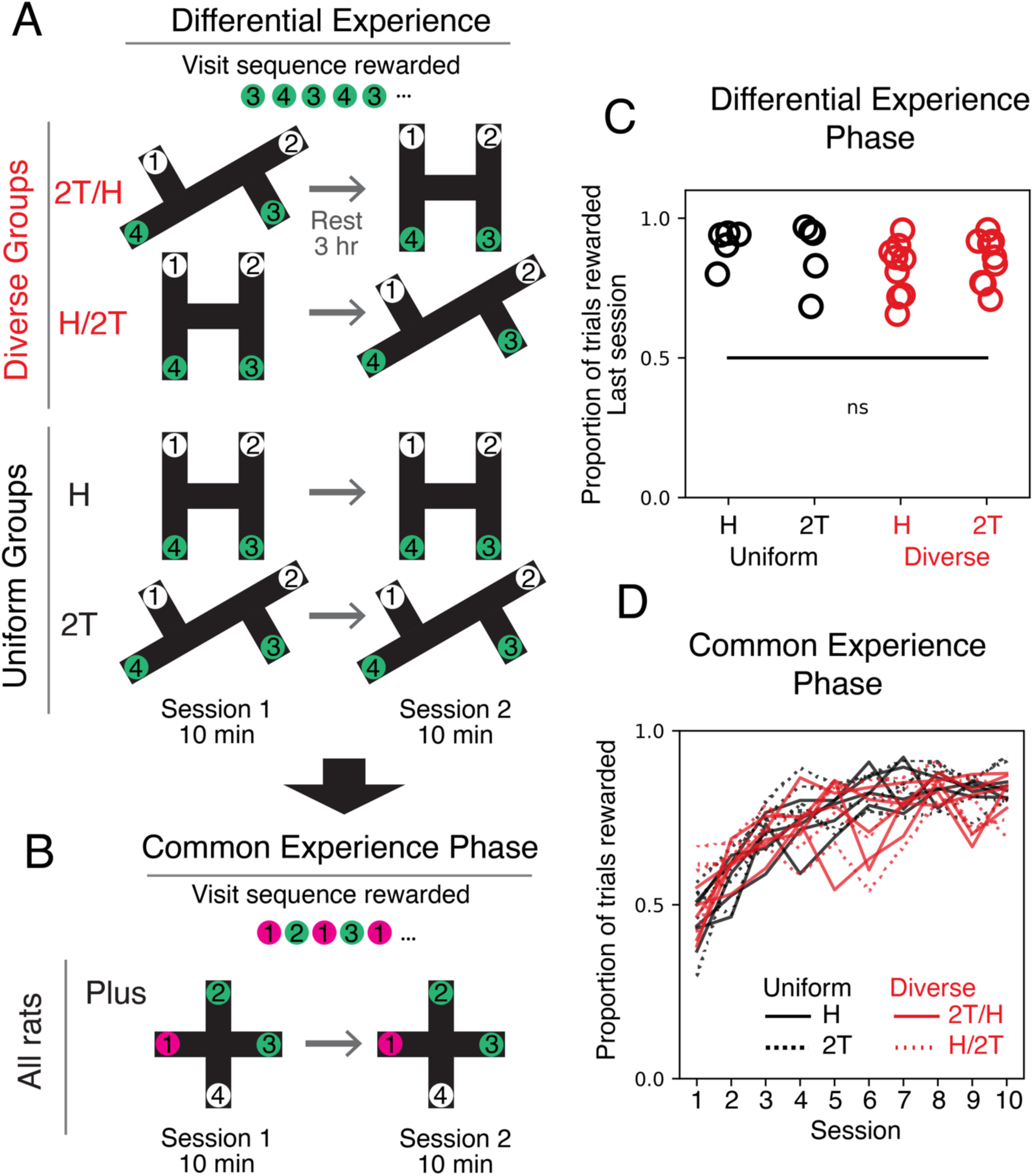
Rats with diverse or uniform experience had similar performance in a novel task. A. Experiment schematic for the differential experience phase. In this phase, all rats were trained for two sessions per day for up to 10 days. Uniform group rats (H and 2T) learned a single task, either the H maze or the 2T maze, and they trained on the same maze during each training session, twice per day. Diverse group (H/2T and 2T/H) rats learned both alternation tasks, H and 2T, counterbalanced for the order of maze sessions. Both the H and 2T mazes have four arm ends. Two of the ends are reward locations (green circles). Visits to the other maze ends (white circles) are not rewarded. The rewarded visit sequence is shown. B. Experiment schematic for the common experience phase. All groups of rats, diverse and uniform, learned to navigate the same maze task for two sessions per day. This was a Plus maze. The rewarded visit sequence is shown. C. Reward rate on the final session of differential experience phase. A two-way ANOVA did not show a significant effect on final performance from experience (p=0.13) or task (p=0.79), or their interaction (F_1, 1_=0.70, p=0.41). D. Proportion of trials rewarded per session on the Plus maze grouped by experience and task in phase 1. Pairwise Wilcoxon rank-sum test with Benjamini/Hochberg false discovery rate correction did not show a significant difference between experience (p>0.44 for all pairs) or task (p>0.83 for all pairs) across sessions.

We asked whether performance on the Plus maze in experiment phase 2 differed between the diverse or uniform experience groups. Based on findings that diverse experiences can benefit cognitive performance (Petrosini et al., 2009), we first predicted that the diverse and uniform groups may differ in performance metrics. Alternatively, given that the maze structure and task rules are sufficiently different between phases 1 and 2 that neither group has any advantages from experience, both groups may show similar performance. We analyzed the reward rate across sessions on the Plus maze and did not find statistically significant differences between experience or task groups (pairwise Wilcoxon rank-sum tests with Benjamini/Hochberg false discovery rate correction: p>0.44 for experience or p>0.83 for task across sessions) (Fig. 1D).

While reward rate did not differ, this metric only quantifies whether the animal’s behavior matches to experimenter-imposed rules and fails to describe the choices made by rats as they learn. Since the tasks required the animals to sequentially visit locations, we hypothesized that the patterns of choices of location visits could provide further insight into the learning process. This may be especially relevant during early learning, when the animals are trying to discover the task rule. We reasoned that the sequence of transitions between consecutive locations may contain patterns at multiple orders (Fig. 2A-B). We defined first-order behavior as the sequence of locations visits. For example, visiting locations 1 and then 2 would comprise a first-order sequence of length 2. Second-order behavior is the egocentric action required to travel between locations: left turn (L), straight (S), or right turn (R). One unit of third-order behavior is created by a pair of two actions, such as a left turn followed by another left turn. We further classified these action pairs into two categories: “similar” corresponds to two turns in a row (for example, R->L) or two straight actions (S->S), and a “dissimilar” action pair, or switch trial, contains a turn and a straight action (for example, R->S or S->L). These categorizations were useful since all mazes in both phases of the experiment had intersections allowing for turns and/or going straight. In contrast, spatial features, such as rewarded locations, were not directly comparable since tasks across experiment differed in topology, geometry, and task rule.

**Figure 2.**
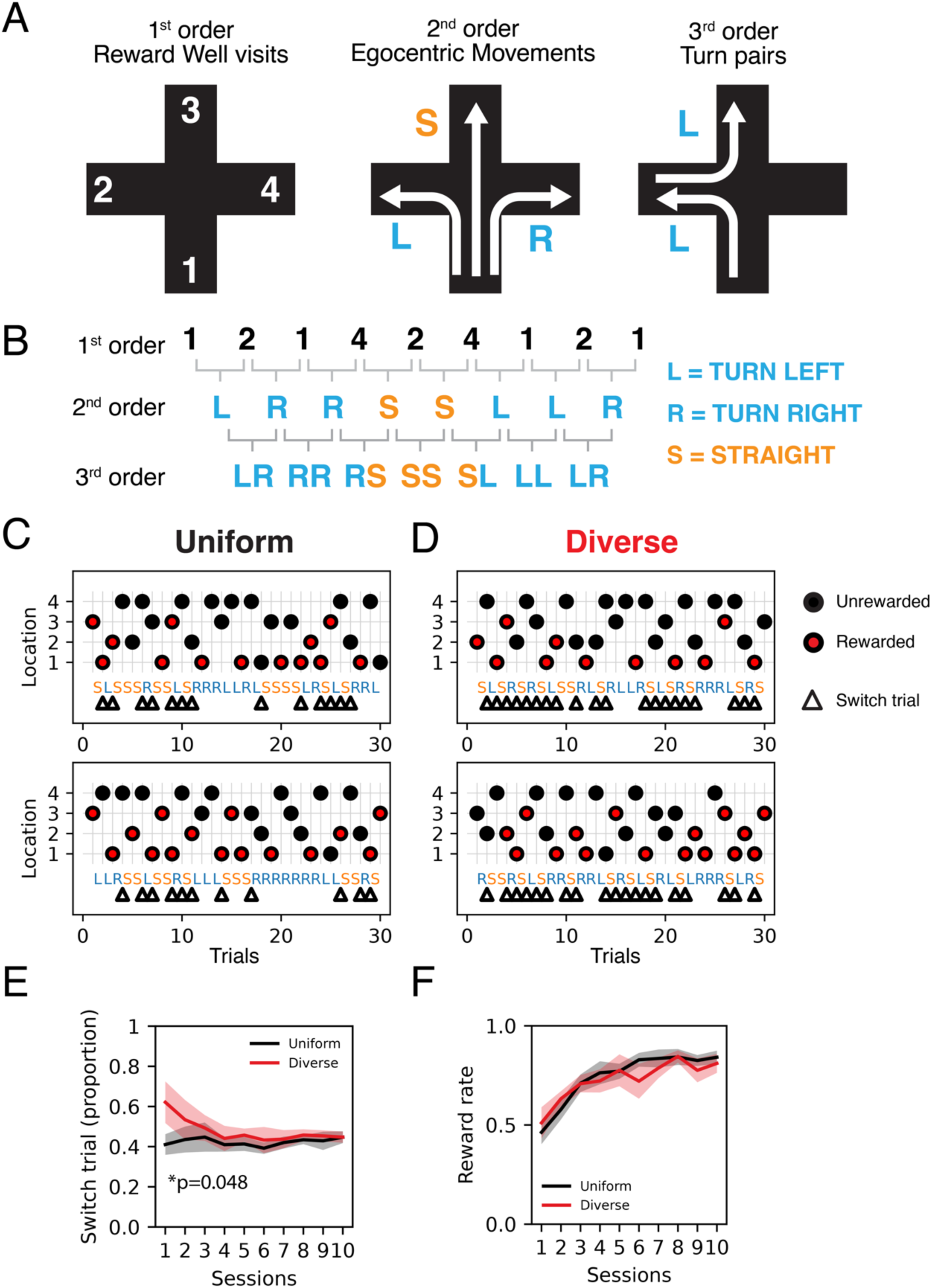
Behavior choice pattern classification for the Plus maze. A. Schematic of 1^st^, 2^nd^ and 3^rd^ order description of behavior choices, which are individual location visits, egocentric movements at junctions, and pairs of turns, respectively. B. Example behavior choice sequence and corresponding higher order descriptions. C. Example behavior choices for the first 30 trials in the Plus maze for 2 animals in the uniform group. 1^st^ order transitions shown by the circles that indicate the maze location visited by the rat. Red circles indicate the rewarded visits. 2^nd^ order transitions convert the location visit pairs into left turns (L), right turns (R) and straight (S). L and R are marked blue and S is in orange. Triangles correspond to switch trials, or 3^rd^ order transitions that involve changes between L/R and S. D. Example behavior choices for the first 30 trials in the Plus maze for 2 animals in the diverse group. E. Proportion of switch trials (mean and 95% confidence interval of the mean) for each session. These are the frequencies of the trial pairs marked by triangles in C and D. Wilcoxon rank-sum test with Benjamin/Hochberg false discovery rate correction, p=0.048 for the first session only. F. Reward rate for each session (mean and 95% confidence interval of the mean) for each session. Data from Fig 1D. is plotted by experience group. Pairwise Wilcoxon rank-sum test with Benjamini/Hochberg false discovery rate correction did not show a significant difference between experience (p>0.44 for all pairs).

Inspection of the raw results shows animals in the diverse group made switch trials more frequently (Fig. 2C-D, Supp. Fig. 1, switch trials indicated with triangles). The proportion of switch trials was greater in the diverse experience group than in the uniform experience group (Fig. 2E), despite both groups having similar reward rates (Fig. 2F). Interestingly, the difference was significant only for the first session (Fig. 2E). This indicates that the diverse experience group were potentially more varied in their choices, switching between turning and not turning at the intersection on consecutive trials, during early learning. These differences disappeared once the animals learned the new rule in later sessions.

The difference in switch trials raised the possibility that the two experience groups made different sequences of choices during phase 2 training. To gain further insight on each animal’s choice sequence, we calculated the probability of each choice sequences for all possible 3-trial sequences. Given sequential choice probability arrays are difficult to visually inspect, we found the dendrogram provides an intuitive visualization (Fig. 3A, Supp. Fig. 2-3), where each node represents a choice, and the connected nodes are subsequent choices (Fig. 3B). The thickness of the edge connecting two nodes indicates the probability of that choice, where thicker lines indicate higher probability (see Methods).

**Figure 3.**
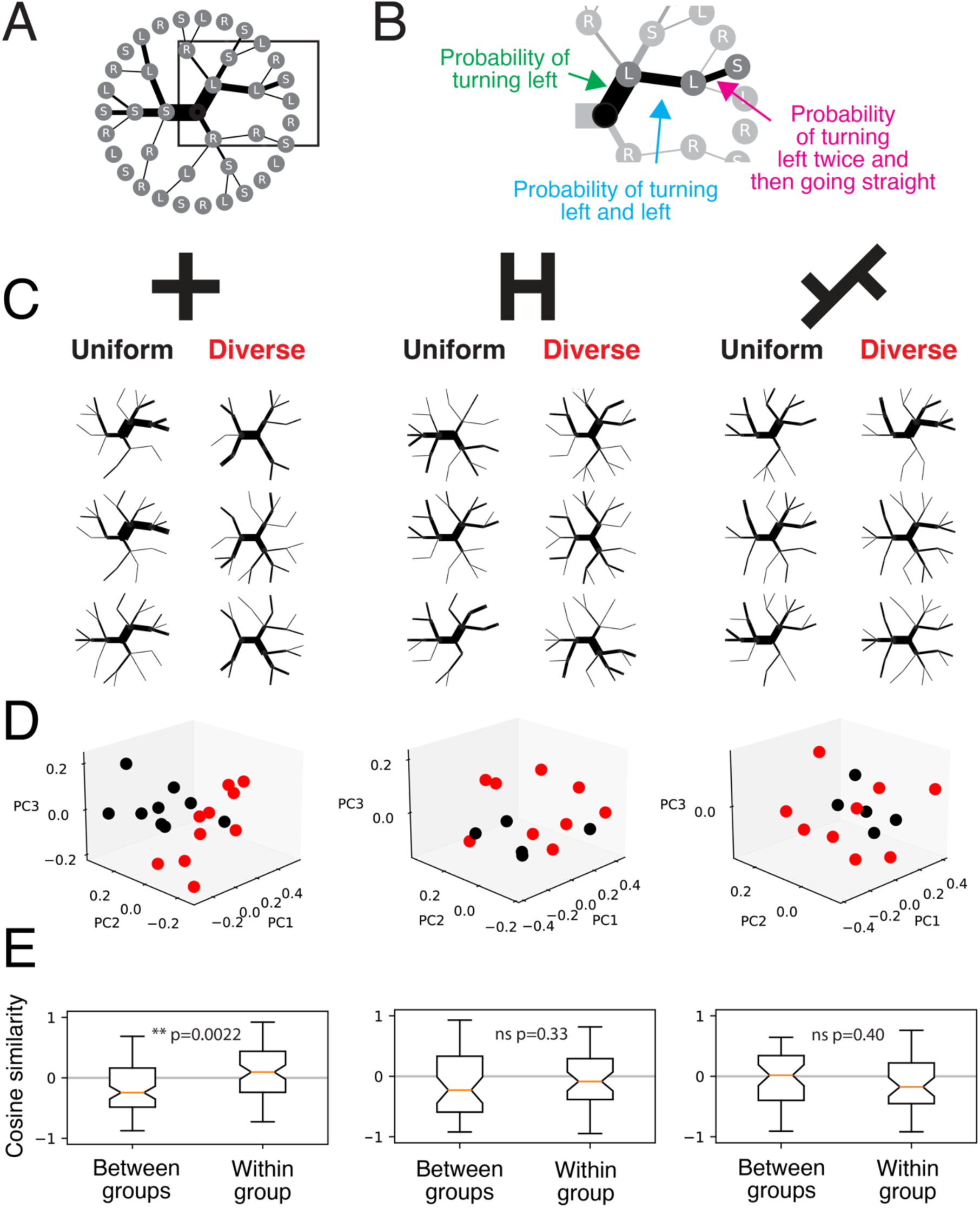
Visualizing sequential behavior choice probabilities. A. Example dendrogram of conditional probabilities for 3-trial choice sequences. Edges represent the conditional probability and nodes represent the choices. B. Example showing the probabilities of sequences with one, two or three trials. C. Choice probability dendrograms for the first 30 trials of the Plus, H and 2T mazes. Three example animals from the uniform (left column) and diverse (right column) are shown. D. Scatter of the first three principal components of the choice probabilities between uniform (black) and diverse (red) groups for the first 30 trials on each maze. To improve visualization of overlapping points, a small jitter (Gaussian noise with standard deviation = 0.003) was added to the values. E. Cosine similarity across all principal components of choice probabilities for between and within groups for the first 30 trials. Wilcoxon rank sum test for Plus (p=0.0022), H (p=0.33) and 2T (p=0.40).

The sequential choice probability dendrograms for the first 30 trials, corresponding to early learning in the Plus maze, revealed distinct patterns within and between groups with uniform or diverse experience in phase 1 (Fig. 3C). To visualize and quantify differences between patterns across the two experience groups, we applied Principal Component Analysis (PCA) on the sequential choice probability array (Fig. 3D). The scatter plots of the first three principal components showed a separation between the uniform and diverse experience groups. We confirmed this separation by calculating the cosine similarity between the principal components of rats. We found that the distance between the uniform experience rats and diverse experience rats was greater than the distance between rats within each experience group, uniform or diverse (Fig. 3E). This is consistent with the idea that uniform and diverse experience groups differ in choice sequence patterns, and we have shown this difference both by the distinct branch structures in the probability dendrograms and by the cosine similarities of the principal components. Dendrograms. We next asked if these differences were experience-dependent or reflected preexisting biases in behavior choices (Kastner et al., 2022). We ruled out the latter explanation since the choice patterns for the first 30 trials on the H or 2T mazes in phase 1 did not differ between the groups (Fig. 3C). Using the same set of analyses, we confirmed all rats had similar behavior choices towards the end of their training when the rule had been learned (Fig. 4, Supp. Fig. 3), consistent with the rats performing the task rule to achieve similar reward rates (Fig. 1C-D).

**Figure 4.**
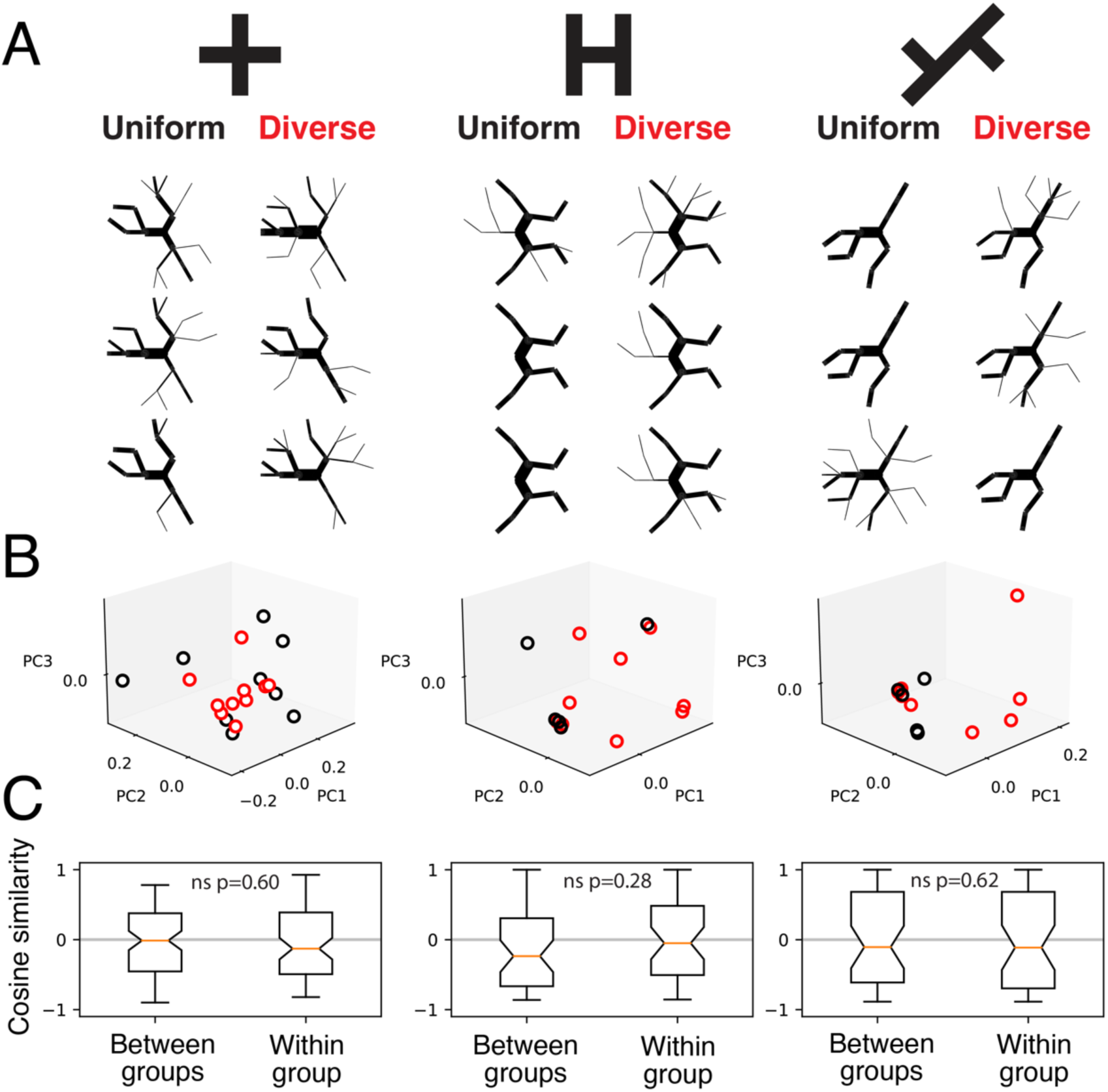
Uniform and diverse groups both converge on the same pattern after training. A. Choice probability dendrograms for the last 30 trials of the Plus, H and 2T mazes. Three animals from the uniform (left column) and diverse (right column) are shown. B. Scatter of the first 3 principal components of the choice probabilities between uniform (black) and diverse (red) groups for the last 30 trials on each maze. To improve visualization of overlapping points, a small jitter (Gaussian noise with standard deviation = 0.003) was added to the values. C. Cosine similarity across all principal components of choice probabilities for between and within groups for the last 30 trials. Wilcoxon rank sum tests for Plus (p=0.60), H (p=0.28) and 2T (p=0.62) mazes.

We next asked what differences in the underlying processes could give rise to these distinct behavior patterns. Our goal is to characterize the structure of the rats’ choice sequences from the early learning period in the Plus maze. We therefore fitted statistical models with a small set of interpretable parameters that relate choice history with future choices using the observed choice sequences for each rat. We then determined whether models from each experience group had different estimated parameter values. Between group differences in model parameters can reveal differences in the statistical processes that generated the sequences. We chose the distance-dependent Chinese Restaurant Process (Blei and Frazier, 2011), which assumes each choice in a sequence is sampled from a distribution of possible choices that is dependent on past choices on two different timescales (Fig. 5A). The model contains a time constant (τ) that determines how influential all past choices are on the next choice, as modeled by an exponential decay. Given that we have found differences in the likelihood of switching between different choices, we added a parameter that determines the relative influence of the previous choice on the next choice (C). The model also included a parameter that determined how closely the probability of choices is biased towards a “base” distribution (ɑ). The base distribution was parameterized to account for simple behavioral strategies based on varying likelihood of repetition (β) without taking more specific trial history into account. Our goal was to compare features of behavioral patterns observed within each experience group during the early learning period by fitting the model to each animal’s behavior during the first 50 trials and comparing the fitted parameters across groups. We performed simulations to show differences in parameters can be recovered from the models (Supp. Fig. 4).

**Figure 5.**
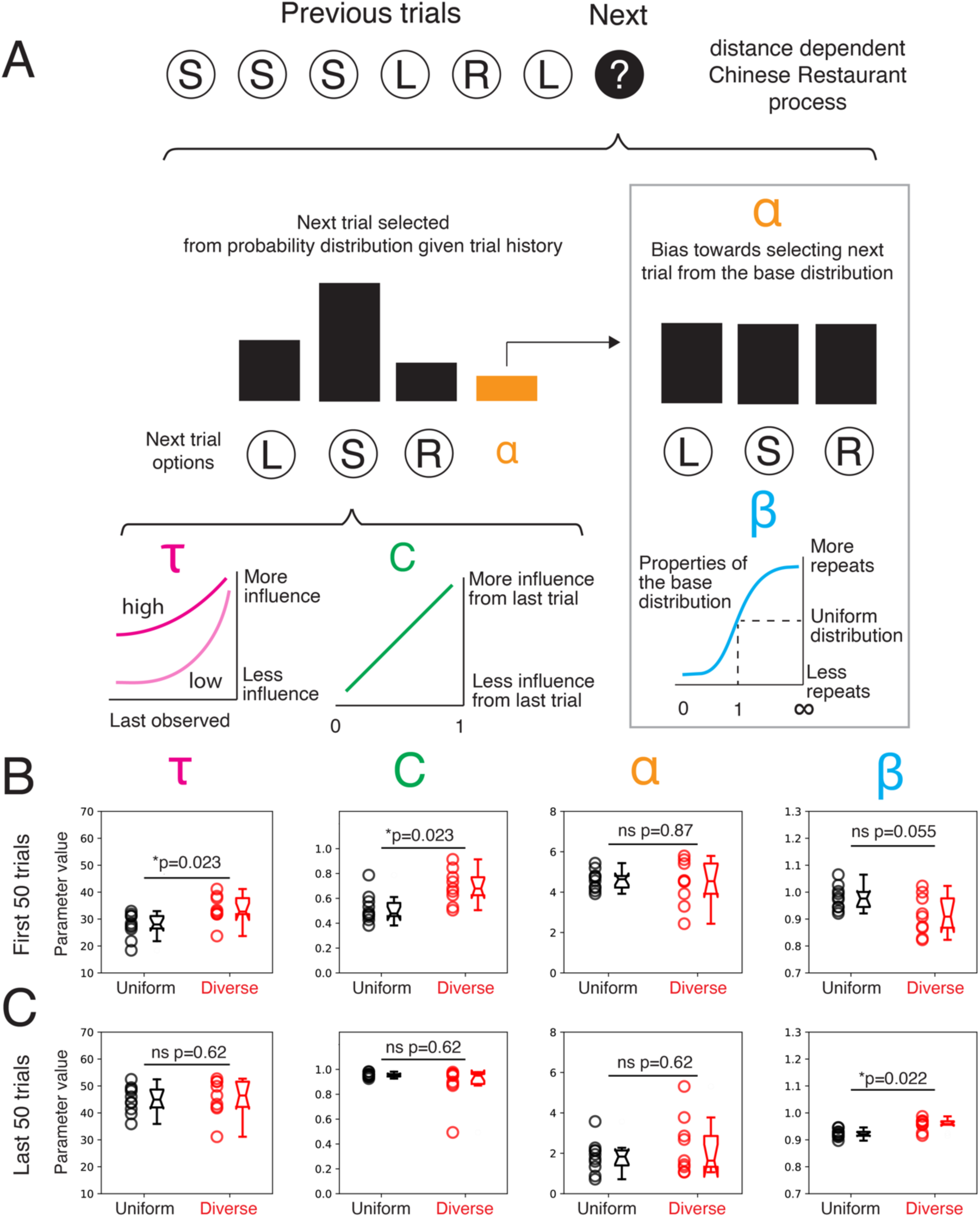
Models of uniform and diverse groups show distinct parameters that control how past trials influence future trials. A. Schematic of a modified distance dependent Chinese Restaurant process model. τ modulates the distance dependent influence of all previous trials on the next trial. C modulates the dependence of the next trial on the immediate previous trial. ɑ determines likelihood the next trial is drawn from a base distribution instead of trial history. β determines the likelihood the base distribution is governed by a uniform distribution or a distribution that is biased to repeat or avoid previous choices. B. Scatter and boxplots of model parameters from model fit to each animal’s first 50 trials on the Plus maze. Wilcoxon rank-sum test with Benjamini/Hochberg false discovery rate correction for τ (p=0.023), C (p=0.023), ɑ (p=0.87) and β (p=0.055). C. Scatter and boxplots of model parameters from model fit to each animal’s last 50 trials on the Plus maze. Wilcoxon rank-sum test with Benjamini/Hochberg false discovery rate correction for τ (p=0.62), C (p=0.62), ɑ (p=0.62) and β (p=0.022).

The model fits showed that the ability of previous choices to predict future choices depended on the experience group. The uniform and diverse groups differed in the estimated time constant (τ), where the diverse group had a significantly longer time constant, indicating that past choices were more predictive of future choices in the diverse group (Fig. 5B). The two groups also differed in how the immediate past affected the upcoming choice (C): the previous choice had a stronger influence for the diverse group of compared with the uniform group. This supports our previous observation that the groups differed in how often they switched between turning and not turning (Fig. 2E). The bias towards selecting the next trial from the base distribution (ɑ) was not significantly different between the two groups. The base distribution was trending to having less repetition (β) in the diverse compared with the uniform group. We confirmed the behavior of both groups converged in the last 50 trials with the model fits producing parameter values (τ, C and ɑ). We found a significant difference in the repetition parameter (β) but the magnitude of the difference is small. This parameter is less influential when the choices are not strongly biased towards the base distribution (ɑ), which is observed for both groups.

Given both statistical and modelling approaches identified differences in choice patterns, we next identified the specific difference in how they switched between choices. Since the diverse experience group had a higher proportion of switch trials at the intersection compared with the uniform experience group (Fig. 2E), we asked whether this can be explained by a higher preference to choose actions not taken previously, an example of which could be turning after going straight on the previous trial or vice versa. Surprisingly, we found the two experience groups differed in the likelihood of switching actions after turning but not after going straight (Fig. 6A). The diverse experience group was more likely to go straight after turning compared with the uniform experience group. Both groups had similar likelihood to turn after going straight. This difference was specific to early exploration in the Plus maze. All groups had similar transition probabilities for the H and 2T mazes and comparable switching likelihoods in the last 50 trials of all mazes, during which behavior across both groups converged (Fig. 6B). We next confirmed this difference was experience-dependent rather than a preexisting difference between the groups. For the animal’s first exposure to the 2T or H mazes, the switching likelihood was similar between groups (Fig. 6A). Further, we found the switching likelihoods in the first or last sessions of the H or 2T mazes was not correlated with those in the Plus maze (Fig. 6C-F). These results indicate different experiences can lead to different specific exploratory choices during searching for solutions to a new problem.

**Figure 6.**
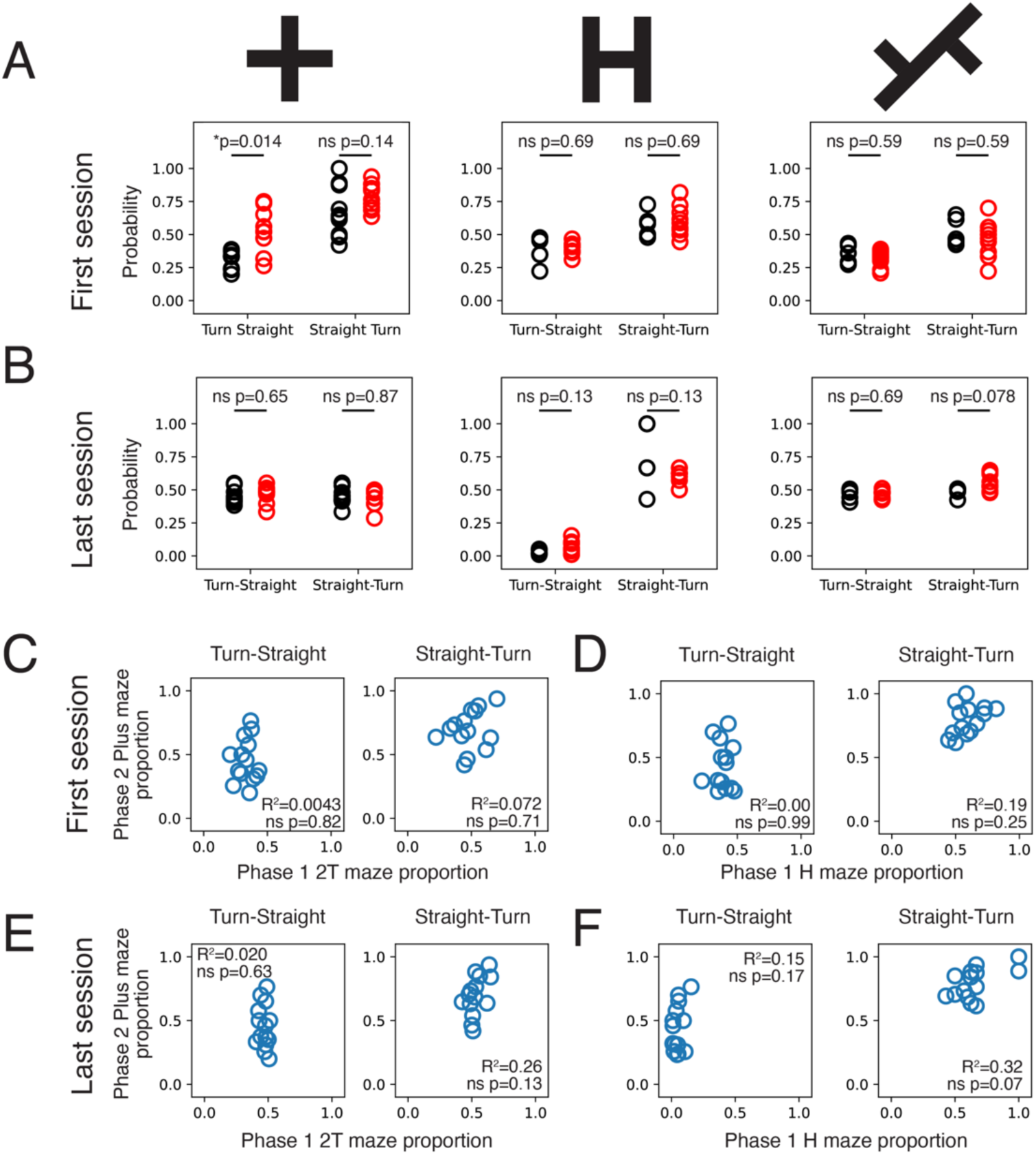
Uniform and diverse groups show distinct likelihood of action switching. A. Probability for turn-straight and straight-turn transitions for the first session on the Plus, H and 2T mazes. Wilcoxon rank-sum p values with Benjamini/Hochberg false discovery rate correction are shown. B. Probability for turn-straight and straight-turn transitions for the last session on the Plus, H and 2T mazes. Wilcoxon rank-sum p values with Benjamini/Hochberg false discovery rate correction are shown. C. Scatter of the proportion of turn-straight and straight-turn transitions across all trials for the first sessions on 2T maze and the first session on the Plus maze. D. Scatter of the proportion of turn-straight and straight-turn transitions across all trials for the first sessions on H maze and the first session on the Plus maze. E. Scatter of the proportion of turn-straight and straight-turn transitions across all trials for the last sessions on 2T maze and the first session on the Plus maze. F. Scatter of the proportion of turn-straight and straight-turn transitions across all trials for the last sessions on H maze and the first session on the Plus maze. All p values for C-F are corrected values using the Benjamini/Hochberg false discovery rate correction.

## Discussion

There are multiple mechanisms through which past experience influences future behavior. Memory is a vessel through which relevant knowledge from one experience can be applied to another, and we can quantify the impact of memory through tests of recall. Generalization involves integrating the memories of multiple experiences to form a broader representation of the concept shared between those experiences, and it is quantified by measuring an individual’s approach to a novel situation that shares attributes, but is not identical, to previous experiences. In the case of multiple problems that share rules or layouts, memory and generalization can allow for the application of previously learned rules, allowing the new problem to be solved faster (Webb, 1917). Our findings reveal that animals’ experience histories can influence their exploratory behavior in a new task. However, the tasks in our experiment did not share spatial topology or navigation rules, removing the utility of directly applying learned spatial layouts or tasks rules from phase 1 to the phase 2. We observed differences in navigation choices during exploration, not performance enhancement. Given this, what could explain the differences in choices made during early exploration in phase 2?

Despite the topological and rule differences between the maze tasks in phases 1 and 2, all mazes have junctions where the rat must decide whether to turn. We hypothesize that the junctions represent a feature shared between mazes, allowing the rats to form abstractions that carry to future tasks. The H, 2T, and Plus mazes all require the rats to make sequences of spatial trajectory choices at junctions, and a sequential turn-based strategy could emerge as they explore and learn. The previous adoption of such a strategy could influence how animals initially explore a new environment that shares the concept of trajectory choices at junctions, even when past spatial geometry and topology are not informative. The turn-based strategy reinforced in the H maze is repeated turns, while turning and going straight is most advantageous in the 2T maze. In phase 2, direct application of these exact sequences of previously learned turns are not possible given the topology of the new maze. However, we hypothesize that previously learned strategies can still bias the likelihood of making certain sequential decisions at junctions in future tasks. This may be particularly applicable to exploration during early learning, before the cognitive map of the new maze has developed. We observed this initial impact through the diverse experience rats’ higher switch trial frequency in the first session.

We hypothesize that experience changed the rats’ approach to exploration throughout the experiment as they were introduced to and learned different tasks. When rats were first introduced to the H and 2T mazes in phase 1, there was variability in choice patterns, which is expected given that rats show preexisting biases in spatial exploration (Kastner et al., 2022). At the end of phase 1, rewarded choice patterns for each task had been reinforced for each animal, and its behavior converged. Their experience in phase 1 determined how they approached exploring a new maze at the start of phase 2. Then, as they learned the new task, their behavior was constrained by the new phase 2 rule, and the behavior across groups converged again.

An important direction for future research is to understand the experience-dependent neural processes that shape learning when neither the application nor the generalization of memory is beneficial or possible. Nonetheless, the animals showed experience-dependent differences in behavior in a novel task. Recent findings show that neural representations of decision-making tasks are dependent on experience (Latimer and Freedman, 2023). Animals with different training histories have distinct cortical task dynamics even when performing the same task with similar behavior. Frontal cortical and hippocampal networks are implicated in representing task rules and modulating generalization (Winocur and Moscovitch, 1990; Freedman, 2001; Wallis et al., 2001; Rich and Shapiro, 2009; Tse et al., 2011; Wang et al., 2012; Xu and Südhof, 2013; Morrissey et al., 2017; Yu et al., 2018; Kaefer et al., 2020; Samborska et al., 2022). These networks could modulate strategic exploratory decisions when learning to solve new problems. When previously learned rules are not applicable, these networks could discard the application of existing rules in favor of more flexible choice policies (Karlsson et al., 2012; Tervo et al., 2014). Experience-dependent changes to neural networks enable the brain to retain information from the past. At any moment, the state of the network is a product of the individual’s unique experience history. We hypothesize that the experience-dependent configuration of the network provides priors for generating new behaviors with a range unique to each individual.

In our experiment, different experience histories led to distinct exploratory patterns in a novel task. Our results suggest that the legacy of past experiences extends beyond the recall of specific memories or the direct application of previously learned concepts. Instead, unique experience histories create unique starting points for how individuals approach new situations. Ours findings challenge us to consider the mechanisms through which experience shapes the differences between each individual and to look beyond memory and generalization for the vessels of transfer between experiences.

## Author Contributions

JYY, NZ and ZML designed the experiment. NZ and ZML collected the data. MPA, JYY and ZS performed the statistical analyses. KWL and JYY performed the computational modelling. MPA, KWL, NZ, ZML and JYY wrote the manuscript.

## Acknowledgements

This work was supported by the University of Chicago Center for Data and Computing Summer Lab program (MPA, ZS) and a Brain Research Foundation Seed Grant BRFSG-2021-11 (JYY).

## Methods

### Pre training

A total of 19 male Long-Evans rats (14∼16 weeks old, 475∼525g) were used in this study. All procedures were performed under approval by the Institute Animal Care and Use Committee at the University of Chicago, according to the guidelines of the Association for Assessment and Accreditation of Laboratory. Animals were kept in a temperature (21°C) and humidity (50%) controlled colony room on a 12/12 h light/dark cycle (lights were on from 8:00 to 20:00). Experiments were performed during the light period. Animals were handled over 4 weeks for habituation to human interaction. Animals were familiarized to foraging for evaporated milk (Carnation) with 5% added sucrose in an elevated black open field box (H: 31cm, W: 61cm, L: 61cm), 10 minutes per day for 3 days. Reward was randomly dropped inside the open box to encourage foraging. Animals were then food restricted to 90% of their baseline weight for 3 days and trained to run back and forth on an elevated linear track (H: 76cm, W: 8cm, L: 60cm) to consume reward from the ends of the track. Animals were trained for 10 minutes per day until a performance criterion of 20 rewards per session (for 4 to 8 days).

### Behavior training

Animals were food restricted to maintain above 85% of their baseline weight. Behavior training was conducted on custom built mazes with interconnecting acrylic track sections (8cm in width) elevated 76cm from the floor. Reward was delivered by a syringe pump (100μl at 20ml/min, NE-500, New Era Pump Systems Inc, New York, USA). Behavior data was recorded using the SpikeGadgets data acquisition system (SpikeGadgets LLC, California, USA). Our experiment was split into two phases. Rats first were exposed to a differential experience phase and then the common experience phase. The common experience phase started 2 days after the end of the differential experience phase.

### Differential experience phase

Animals were randomly assigned to the uniform (n=10) and diverse group (n=9). A H-maze and a double T-maze (2T) were used for the differential experience phase training (Figure 1A). Animals obtained reward from the ends of the maze only when visiting two of the four ends in alternation. Animals performed two 10-minute training sessions separated by 3 hours, during which the rats were returned to their cages. Training continued until performance exceeded 80% or up to 10 days. The diverse group trained on two mazes per day, while the uniform group trained on one maze twice per day. For the diverse group, we controlled in the order the rats learned the tasks across each day by assigning 5 rats to 2T maze first then H maze. The remaining 4 rats learned the task in reverse order.

### Common experience phase

After the differential experience phase, all animals (n=19) were trained on a Plus maze with a different rule (Fig. 1B). Both groups were trained to visit three of the four wells in a specific sequence to receive reward at those three wells (Fig. 1B). The sequence involves alternating visits between well 1 and wells 2 and 3. Animals underwent two 10-minute training sessions per day, for 5 days. The rats were returned to their cages for 3 hours between the two sessions.

### Data processing and analysis

We registered reward well visits based on sensor trigger events and reward delivery based on pump trigger events. All analyses were performed in Python using Numpy, Scipy and scikit-learn.

### Behavior pattern classification

We started with a sequence of reward location visits, which represent first-order patterns. We converted this sequence into second-order behavior patterns given each pair of transitions requires one to two movement choices: left turn (L), straight (S), or right turn (R). We then classified third-order patterns as the transition between 2^nd^ order actions, such as a left turn followed by another left turn. We can further classify these action pairs into similar, corresponding to turn followed by turn or straight followed by straight, versus dissimilar, or a switch trial, corresponding to turn followed by straight or vice versa.

### Choice sequence probabilities

We calculated the probability of observing a specific choice sequence for all possible 3-trial sequences, for example (left, left left, left left left, …). Data for all rats is in the form of a *m×n* matrix with *m* being each animal and *n* being all the possible sequences. To visualize the probability matrix as a dendrogram, we used the Python networkx package (https://networkx.org/). To visualize the similarity between the probability matrices for the uniform and diverse groups, we then used Principal Component Analysis to reduce the dimensionality of this matrix. To quantify similarity, we calculated the pairwise cosine similarity for a pair of animals across all principal components. This was done for within (diverse to diverse, uniform to uniform) and across (diverse to uniform) group comparisons.

### Modified distance-dependent Chinese restaurant process model

We aimed to summarize statistically how the actions of each rat in the Plus maze depended on the recent trials and how the distribution of choices changed over the course of learning. Given the sequence of trials performed by an animal, we modeled the action on a trial as a probability distribution that depended on the past trial and the number of trials performed. The dependency of the number of trials allowed the model to account for the changes in the animals’ behavior during learning. This contrasts to a typical Markov model, which assumes behavior only depends on the past trials but not the history of trials performed.

To accomplish this, we modeled the sequence of actions (left, right, or straight) performed by each rat using a sequential distance-dependent Chinese restaurant process model (ddCRP) (Blei and Frazier, 2011). We modified the model by adding a parameter that specifically controls the contribution of the last trial to the upcoming choice. The ddCRP defines a generative stochastic process in which the probability of the action on the *i*th trial depends on the outcomes of the previous trials. The probability of observing action A on trial *i*(*y_i_*) is given as:

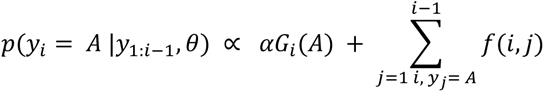

Where the distance function between trials *i* and *j* is

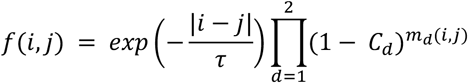

Where *m_d_*(*i*,*j*) = 0 if trials *i* and *j* share the same context of depth *d*: that is, the sequence of *d* actions immediately preceding trials *i* and *j* are the same. Otherwise, we set *m_d_*(*i*,*j*) = 1. The timescale parameter of the distance function, τ > 0, determines how predictive actions from the past are of the current trial. Low values of τ indicate that the actions at the beginning of the session are not informative of the animals’ behavior at the end of the session. This timescale gives the process the “distance dependent” property in comparison to the standard Chinese restaurant process, which weighs all previous observations with weight 1. The context parameters, *C_d_* ε [0,1], determine how much choice depends on specific actions permed on the *d* previous trials (the context). If *C_d_* = 1, then context is weighted heavily by the model: the actions performed in one context do not inform the actions in a different context. If *C_d_* = 0, context is not predictive of the actions.

The remaining two parameters define the *base measure*, *G_i_*: the prior probability over the actions.

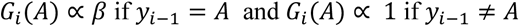

The concentration parameter, *α* > 0, determines bias for selecting the choice on each trial from the base distribution. The bias parameter, *β* > 0, is included to alter the base distribution. The value of this parameter account for how a fixed switch-stay bias could account for the animals’ sequence of actions. Actions are drawn from the uniform distribution *a priori* if *β* = 1. For *β* < 1, actions are less likely to be repeated, and for *β* > 1, choices are more likely to be repeated.

Our approach extends the ddCRP model for sequences to include recent context within the distance function. This approach is inspired by models that use hierarchical Dirichlet priors to regularize estimation of Markov models (Wood, 2009). However, our method takes advantage of the fact that the distance function already weighs the previous observations differently. Thus, we could incorporate dependencies on recent actions without a more complex hierarchical model in contrast to a recently proposed statistical model of behavioral sequences (Éltető et al., 2022).

We fit the model Markov chain Monte Carlo (MCMC) methods in a Bayesian framework implemented using the Stan modeling platform (STAN Development Team, 2023). Convergence of the MCMC procedure was assessed using the *R*^ metric (Vehtari, 2021) with four independent chains of 1000 samples each. We used the posterior median as a point estimate for individual parameters. The prior distributions for the parameters were independent for each parameter:

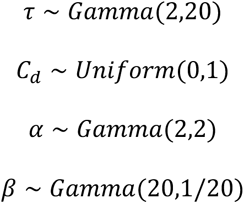

where the gamma distributions are parameterized as shape and scale.

---

**Supplementary Figure 1.**
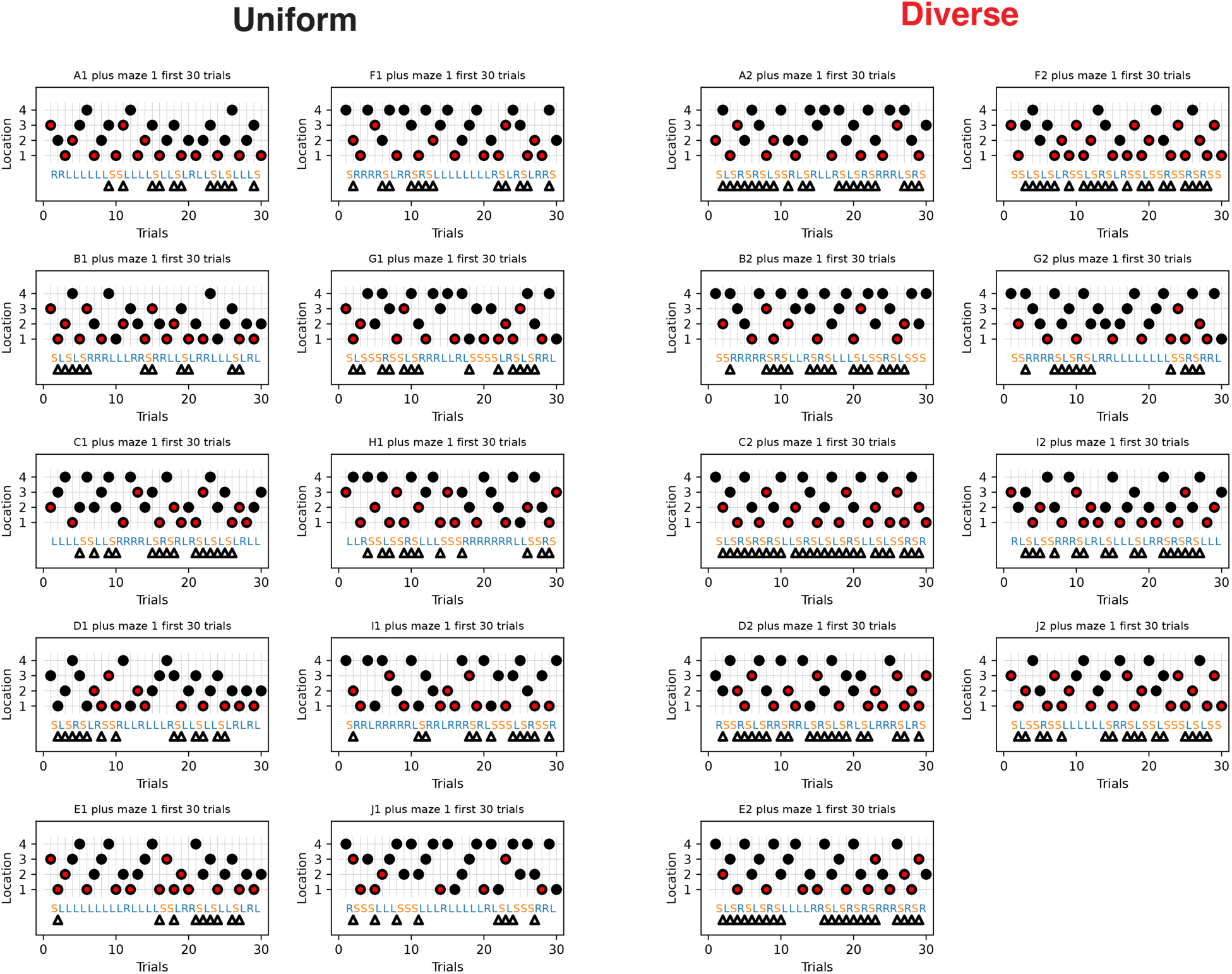
Behavior choices for the first 30 trials of the Plus maze for all animals, shown in the same format as Fig. 2 C-D. 1^st^ order transitions shown by the circles that indicate the maze location visited by the rat. Red circles indicate the rewarded visits. 2^nd^ order transitions convert the location visit pairs into left turns (L), right turns (R) and straight (S). L and R are marked blue and S is in orange. Triangles correspond to switch trials, or 3^rd^ order transitions that involve changes between L/R and S.

**Supplementary Figure 2.**
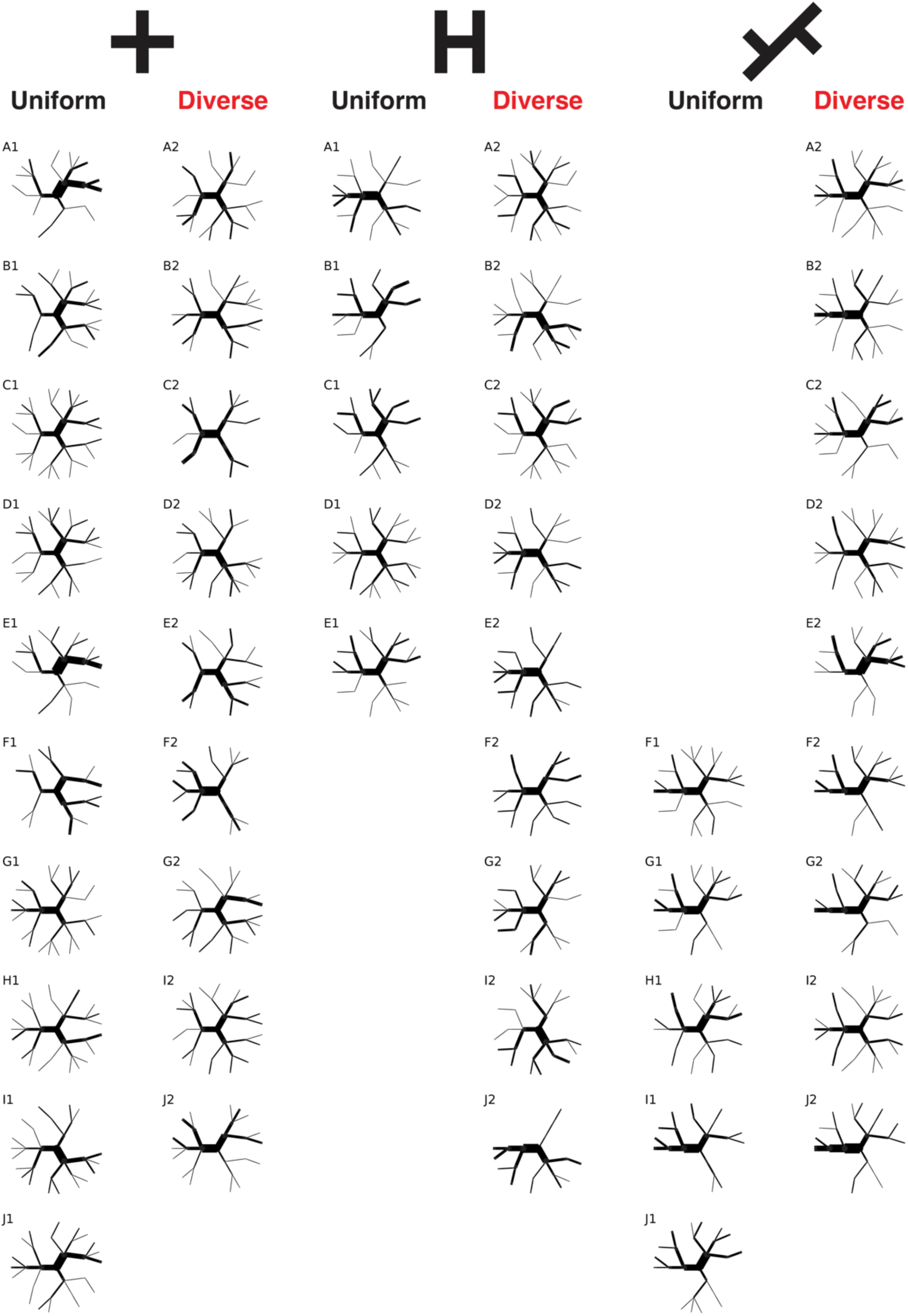
Choice probability dendrograms for the first 30 trials of the Plus, H and 2T mazes for all animals.

**Supplementary Figure 3.**
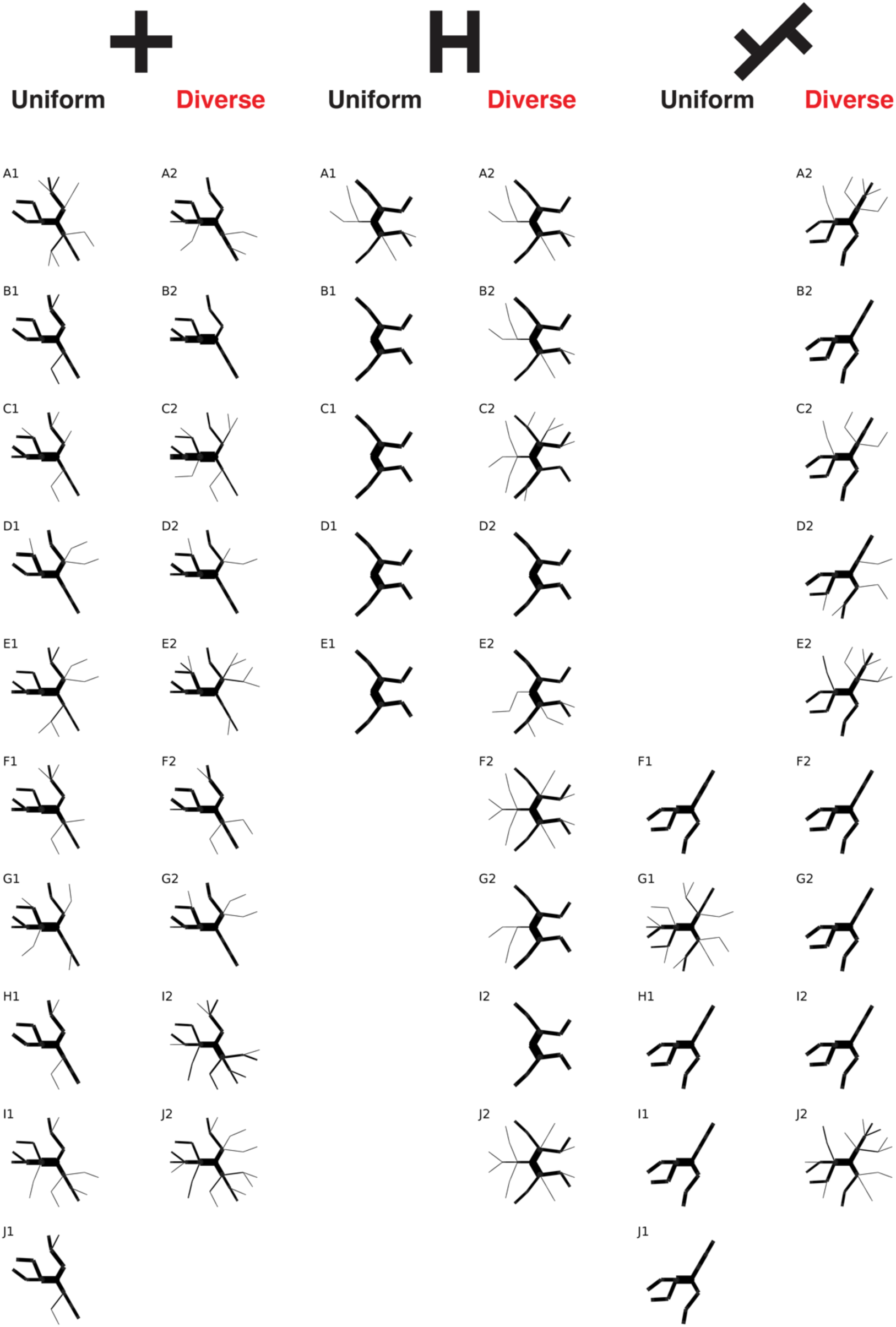
Choice probability dendrograms for the last 30 trials of the Plus, H and 2T mazes for all animals.

**Supplementary Figure 4.**
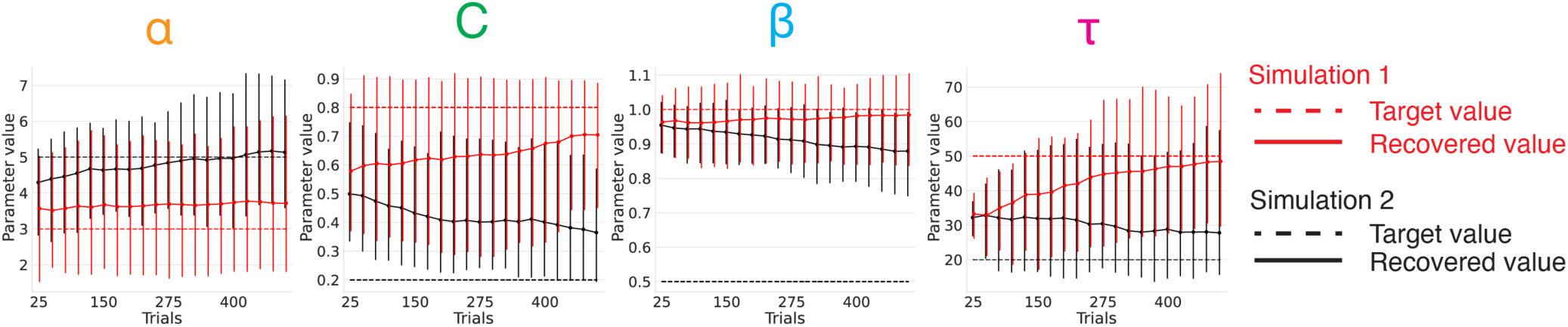
Simulations confirm that the model parameters can be recovered from sequences of actions. We simulated from the distance-dependent Chinese restaurant process using two different sets of parameters (simulations 1 and 2, dashed lines indicate the true parameters). For each set of parameters, we generated 50 independent simulations. The parameters were then fit with an increasing number of trials using the posterior median as the estimate. The points give the mean estimates and the error bars show a 90% interval over simulations. The estimated parameters remained close to the prior distribution with few trials, and tended towards the true parameters with increasing amounts of data. We found that the context dependency parameter (C) required the fewest number of trials to separate across these two simulations. Given low values of the chosen base distribution bias (ɑ), which meant the base distribution was unlikely to be chosen in the generated sequences compared with the history-dependent distributions, we did not expect the repetition bias parameter (β) to be effectively recovered.

## Notes

### Competing Interest Statement

The authors have declared no competing interest.

